# Human iPS cell-derived sensory neurons can be infected by SARS-CoV-2 strain WA1/2020 as well as variants delta and omicron

**DOI:** 10.1101/2023.01.10.523422

**Authors:** Anthony Flamier, Punam Bisht, Alexsia Richards, Danielle Tomasello, Rudolf Jaenisch

## Abstract

COVID-19 has impacted billions of people in the world since 2019 and unfolded a major healthcare crisis. With an increasing number of deaths and the emergence of more transmissible variants, it is crucial to better understand the biology of the disease-causing virus, the SARS-CoV-2. Peripheral neuropathies appeared as a specific COVID-19 symptom occurring at later stages of the disease. In order to understand the impact of SARS-CoV-2 on the peripheral nervous system, we generated human sensory neurons from induced pluripotent stem cells that we infected with the SARS-CoV-2 strain WA1/2020 and the variants delta and omicron. Using single cell RNA sequencing, we found that human sensory neurons can be infected by SARS-CoV-2 but are unable to produce new viruses. Our data suggests that sensory neurons can be infected by the original WA1/2020 strain of SARS-CoV-2 as well as the delta and omicron variants.

## Introduction

Infection with SARS-CoV-2 has been reported to impact the entire body and cause COVID-19^1^. In addition to causing severe damage to the respiratory system, SARS-CoV-2 acutely affects the nervous system with symptoms including loss of taste and smell, headaches, stroke, delirium, and brain inflammation^2,3^. Among these symptoms, anosmia emerged as an early indication of SARS-CoV-2 infection^4^. The observed anosmia was highly prevalent in COVID-19 patients but reversible, with most patients recovering their senses of smell and taste in about 6 weeks. However, in about 10% of COVID-19 patients, olfactory dysfunction becomes either persistent or poorly recovered^4–11^. Recent *in vivo* studies suggested that SARS-CoV-2 can disrupt the nuclear architecture of the olfactory epithelium, inducing the dysregulation of olfactory receptor genes in a non-cell autonomous manner^12^. In this study, using a model of SARS-CoV-2 intranasal injection of hamsters^12^, the authors also noted that sensory neurons of the olfactory epithelium were poorly infected (<5% positivity). Although olfactory sensory neurons appear to not express the receptor for virus entry,^12^ it remains unclear whether other types of peripheral sensory neurons may be susceptible to SARS-CoV-2 infection. Additional COVID-related peripheral neuropathies have been described recently including small-fiber neuropathy, multifocal demyelinating neuropathy and critical illness axonal neuropathy^13,14^. It is estimated that 59% of COVID-19 patients show signs of neuropathy in the mid- or long-term^13^. The available data suggest that neurons are not highly infected by SARS-CoV-2 due to the lack of ACE2 expression and that neuropathies could be due to an inflammatory response affecting sensory neurons in a non-cell-autonomous manner^12–14^. However, neuropathy disorders generally occur at a late stage of the disease when the inflammatory response is dimmed^13,14^.

In order to address the mechanism by which SARS-CoV-2 impacts the peripheral nervous system, we generated a heterogeneous population of human sensory neurons using induced pluripotent stem (iPS) cells that were infected with either the WA1/2020 strain of SARS-CoV-2 or the delta and omicron variants. Single-cell RNA sequencing (scRNAseq) analysis showed that about 20% of human sensory neurons were infected by SARS-CoV-2 with the omicron variant having the lowest infection rate. We further show that although SARS-CoV-2 infects human sensory neurons, it does not actively replicate to shed new viruses. This study reveals how SARS-CoV-2 may initiate a non-productive infection in sensory neurons which has the potential to explain the neuropathies associated with COVID-19.

## Results

### Human sensory neurons express the SARS-CoV-2 receptor gene ACE2

SARS-CoV-2 can access the host cells through the binding of its spike protein to the extracellular domain of ACE2 protein^1^. Cells lacking ACE2 receptor fail to be infected *in vivo* and *in vitro* ^15,16^. Conversely, cells overexpressing ACE2 show a higher rate of infection ^16,17^. To evaluate whether human sensory neurons express ACE2 and could potentially be infected by SARS-CoV-2, we differentiated human iPS cells into sensory neurons using established protocols^18^. After 1.5 months of differentiation, more than 90% of the cells expressed sensory neuron markers such as Trk.B, Trk.C, Ret, *CGRP* and *PRPH* (Figure S1A and data not shown). When compared to other cell types, iPS-derived sensory neurons showed robust expression of *ACE2* transcript similar to lung cells (Calu3) indicating their infectibility (Figure S1B).

### SARS-CoV-2 infects human sensory neurons

Based on the previous observations, it remains unclear whether SARS-CoV-2 may be able to enter human sensory neurons. In order to address this question, we performed scRNAseq on iPS-sensory neurons exposed to heat-inactivated (HI) SARS-CoV-2 or fully active WA1/2020 SARS-CoV-2. We added HEK293T (293T) cells in the culture in order to simultaneously compare iPS-sensory neurons to cells with similar expression of *ACE2* (Figure S1B) replicating actively the virus. The different gene expression patterns in the two cell types allowed to separate two well-defined clusters containing the same proportion of mock, HI, and SARS-CoV-2 exposed cells (Figure 1A). 293T cells were distinguished from neurons by expression of the *AMOT* and *SV40T* genes (Figure 1B). *AMOT* encodes for a protein that controls the spatial distribution of apical polarity proteins in kidney^19^ and is expressed in 293T cells. Another marker to distinguish 293T cells from neurons is expression of SV40 large T antigen used to derive the cells. Notably, we identified a subcluster of more mature sensory neurons expressing high levels of the sensory markers *NTRK2* (coding for Trk.B) and *NTRK3* (coding for Trk.C) (Figure 1B). When comparing cells from each group, we detected SARS-CoV-2 RNA in about 30% of the cells exposed to the virus (Figure S1C). For further analysis, we looked at Nucleocapsid expression in 293T cells (positive for *SV40T*) as well as sensory neurons (positive for *NTRK2* and *NTRK3*). In cells exposed to live SARS-CoV-2, Nucleocapsid RNA was detected in 21% of 293T cells and 18% of sensory neurons, but was not detected in mock and HI-treated cells (Figure 1C and S1C) suggesting that sensory neurons support SARS-CoV-2 infection and RNA replication. We compared gene expression profiles of sensory neurons positive for SARS-CoV-2 RNA with that of SARS-CoV-2 RNA negative sensory neurons. Similar levels of sensory neuron and neural crest cell markers were observed between both populations suggesting that the maturity of sensory neurons is not a determinant of infection. Instead, SARS-CoV-2 RNA positive cells showed a reduced expression for genes coding for RNA metabolism proteins such as *FUS, NUCKS1* and *NCL* (Figure 1D). It has been shown that SARS-CoV-2 remodels the host cell RNA metabolism machinery and inhibition of RNA metabolic pathways can inhibit SARS-CoV-2 infection^20^. Our results suggest that infected sensory neurons reduce their RNA metabolism activity which may inhibit viral RNA translation. Additionally, olfactory receptor genes were not highly expressed in both populations which correlates with previous observations that SARS-CoV-2 does not infect olfactory sensory neurons of COVID-19 patients (data not shown and Figure 1D)^12^. In summary, our data show that SARS-CoV-2 infects human sensory neurons.

**Figure 1:**
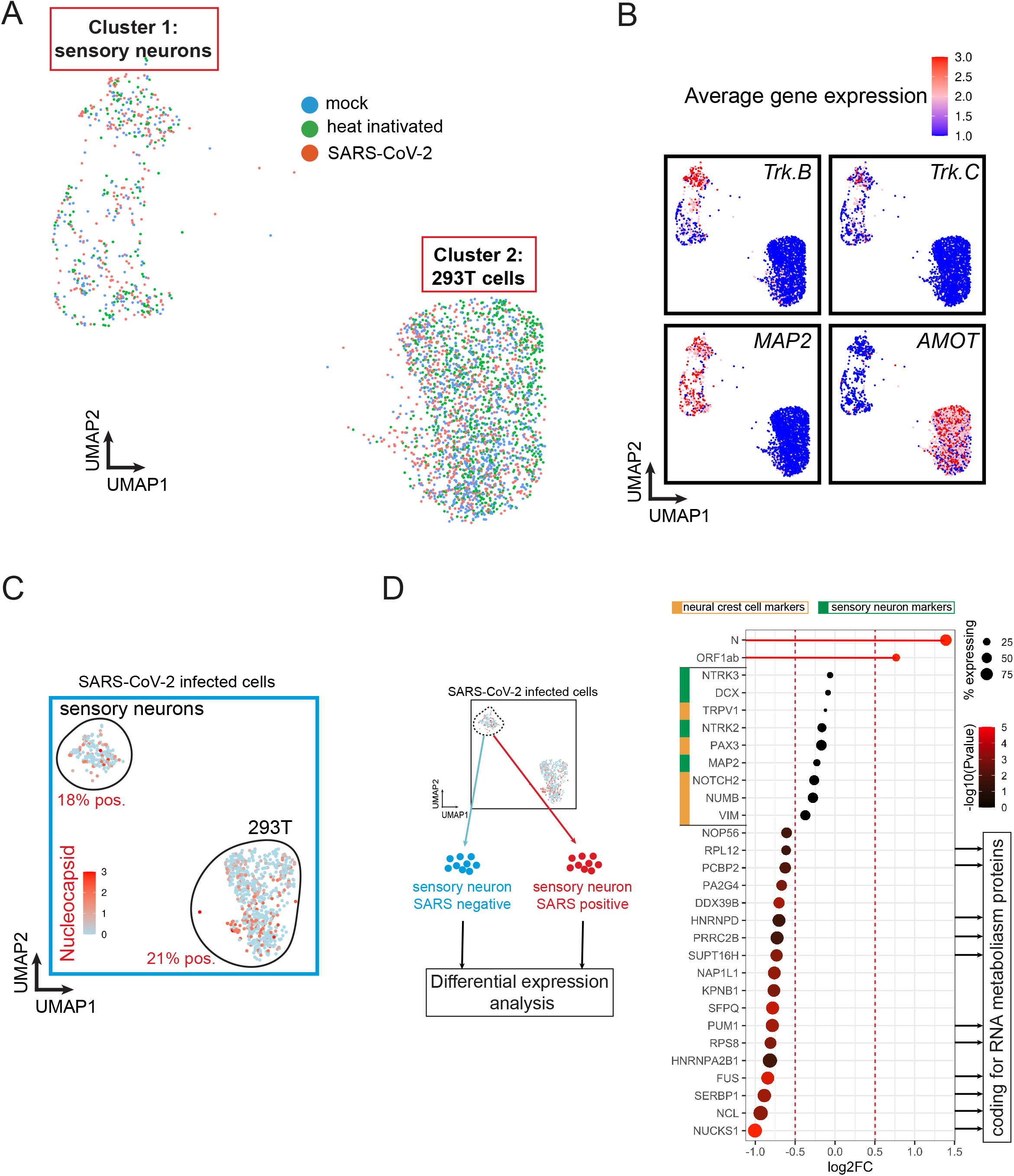
SARS-CoV-2 infects a subset of human sensory neurons. (A) UMAP of single-cell RNA sequencing (scRNAseq) from hiPSC-sensory neurons co-cultured with HEK293T cells exposed to heat-inactivated virus or SARS-CoV-2 strain WA1/2020 for 48h. Mock is used as a control. (B) Feature plot of average gene expression for sensory markers *NTRK2* (Trk.B), *NTRK3* (Trk.C), pan-neuronal marker *MAP2* and 293T cell marker *AMOT* in all samples combined (mock, heat-inactivated and SARS-CoV-2). (C) Feature plot of average gene expression for SARS-CoV-2’s Nucleocapsid in hiPSC-sensory neurons co-cultured with HEK293T cells exposed to SARS-CoV-2 for 48h. (D) Differential gene expression analysis between SARS-CoV-2 positive and negative sensory neurons. The dot size indicates the proportion of sensory neurons expressing the corresponding gene. The color gradient of each dot indicates the P-value for the change in gene expression (more red=more significant). Genes with a negative Log2FC have reduced expression in SARS-CoV-2 positive neurons compared to SARS-CoV-2 negative neurons.

### Infected sensory neurons synthesize SARS-CoV-2 RNA but do not produce new viruses

The SARS-CoV-2 genome is a positive sense, single stranded RNA. Upon entry into the host cell, SARS-CoV-2 uses the host cell machinery to initially generate a complementary negative sense genome length RNA. The negative strand is used as a template for the production of additional positive strand genomes which serve both as a template for translation of viral proteins and are incorporated into nascent virions^21^. The presence of negative-sense SARS-CoV-2 RNA can therefore be used as a marker of SARS-CoV-2 RNA replication. To confirm that SARS-CoV-2 RNA is replicating in both sensory neurons and 293T cells, we examined scRNAseq data for the presence of negative strands of SARS-CoV-2 RNA. Negative strands of SARS-CoV-2 RNA were detected in 293T cells infected by all three of the variants tested suggesting active viral RNA replication in these cells (Figure 2A). The proportions of negative strands compared to positive strands was similar to what was previously observed in infected cell lines or COVID patient samples^22^. In sensory neurons, negative strand RNA was detectable only in WA1/2020 and delta exposed cells (Figure 2A). This indicates that SARS-CoV-2 RNA is actively synthesized in sensory neurons upon infection with WA1/2020 SARS-CoV-2 as well as the variant delta. Sensory neurons infected by the omicron variant did not show any sign of viral RNA synthesis at this timepoint. It is also worth noting that the proportion of negative strand SARS-CoV-2 RNA compared to the positive strand is higher in sensory neurons exposed to the delta variant as compared to the WA1/2020 strain (Figure 2A). This difference could be explained by either a difference in the rate of infection or in the rate of viral RNA replication.

**Figure 2:**
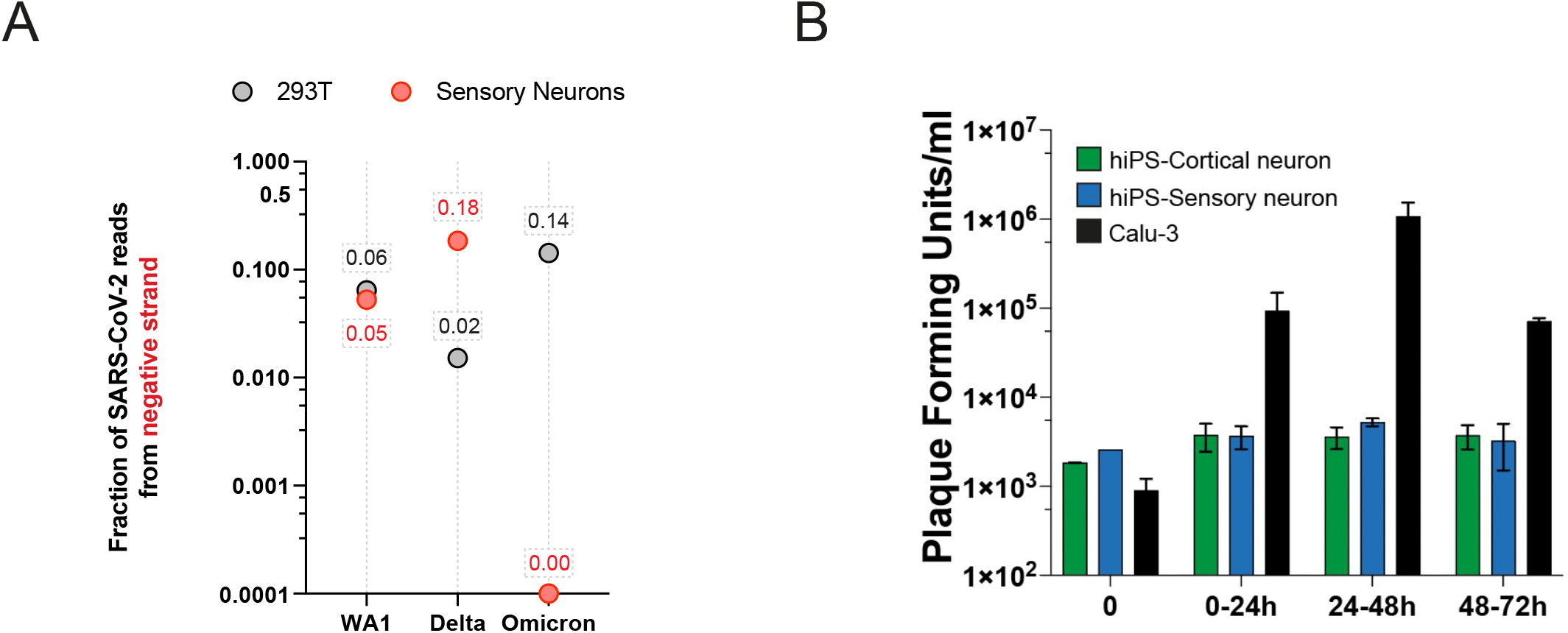
Infected sensory neurons synthesize SARS-CoV-2 RNA but do not replicate new viruses. (A) Fraction of negative strand SARS-CoV-2 RNA compared to the total number of SARS-CoV-2 RNA. Negative strand SARS-CoV-2 RNA was detected in sensory neurons and 293T cells positive for SARS-CoV-2 RNA in scRNAseq. (B) Plaque forming assay on hiPSC-derived cortical and sensory neurons and Calu-3 cells exposed to WA1/2020 SARS-CoV-2 for 24, 48 and 72 hours. N=3; t-test *p-value<0.05. Mean +/- S.E.M.

To determine if sensory neurons could shed infectious SARS-CoV-2 we infected the cells with the virus and quantitated the amount of infectious virus in the media by plaque assay. The lung carcinoma line Calu-3 was used as a positive control. At 48 hours post infection the amount of infectious virus in the media was similar to cells not expressing ACE2 such as hiPS-cortical neurons (Figure 2B). Conversely, Calu-3 cells showed robust amplification of infectious virus by 48 hours post infection without any sign of cytotoxicity (Figure 2B). These observations suggest that human sensory neurons do not propagate SARS-CoV-2 *in vitro*. In summary, these data highlight that sensory neurons exposed to SARS-CoV-2 replicate SARS-CoV-2 RNA but do not shed new virus. The mechanism by which the sensory neurons block the assembly and release of new viruses remains to be elucidated.

### Infection of sensory neurons by SARS-CoV-2 variants delta and omicron

Peripheral neuropathies have been reported in many COVID-19 cases throughout the world^13,14,23^. The most common form of peripheral neuropathy associated with SARS-CoV-2 infection is anosmia, with cases reported from the first wave of the pandemic when the variant alpha was predominant^13,14,23^. While the delta variant seemed to equally affect the peripheral nervous system, the more transmissible and less severe omicron variant showed a decrease in reported anosmia and peripheral neuropathies^24^. To assess differences in the infection of sensory neurons by WA1/2020 SARS-CoV-2 and the delta and omicron variants we infected co-cultures of iPS-derived sensory neurons and 293T cells for 48h and analyzed the cells by scRNAseq. Cells positive for SARS-CoV-2 RNA were detected in both sensory neurons and 293T cells for the three variants (Figure 3A-B). The delta variant was able to infect more 293T cells than the WA1/2020 strain (32% compared to 17%) but less sensory neurons (21% compared to 25%) as determined by Nucleocapsid RNA levels (Figure 3A-B and S2A-B). Conversely, the omicron variant infected three times less sensory neurons and 293T cells than the WA1/2020 strain and delta variant which is consistent with our data and observations from the literature with the omicron variant having a slower rate of propagation at the beginning of the infection in some cell types^25^ (Figure 3A-B and S2A-B). Notably, while Nucleocapsid and Orf1ab RNA levels were the most abundant transcripts, other SARS-CoV-2 genes such as the Spike (S) and the Membrane (M) could be detected in both sensory neurons and 293T cells (Figure S2C). These results show that SARS-CoV-2 strain WA1/2020 and variants delta and omicron may have different tropisms and rates of infection for sensory neurons but they are all able to infect both sensory neurons and 293T cells.

**Figure 3:**
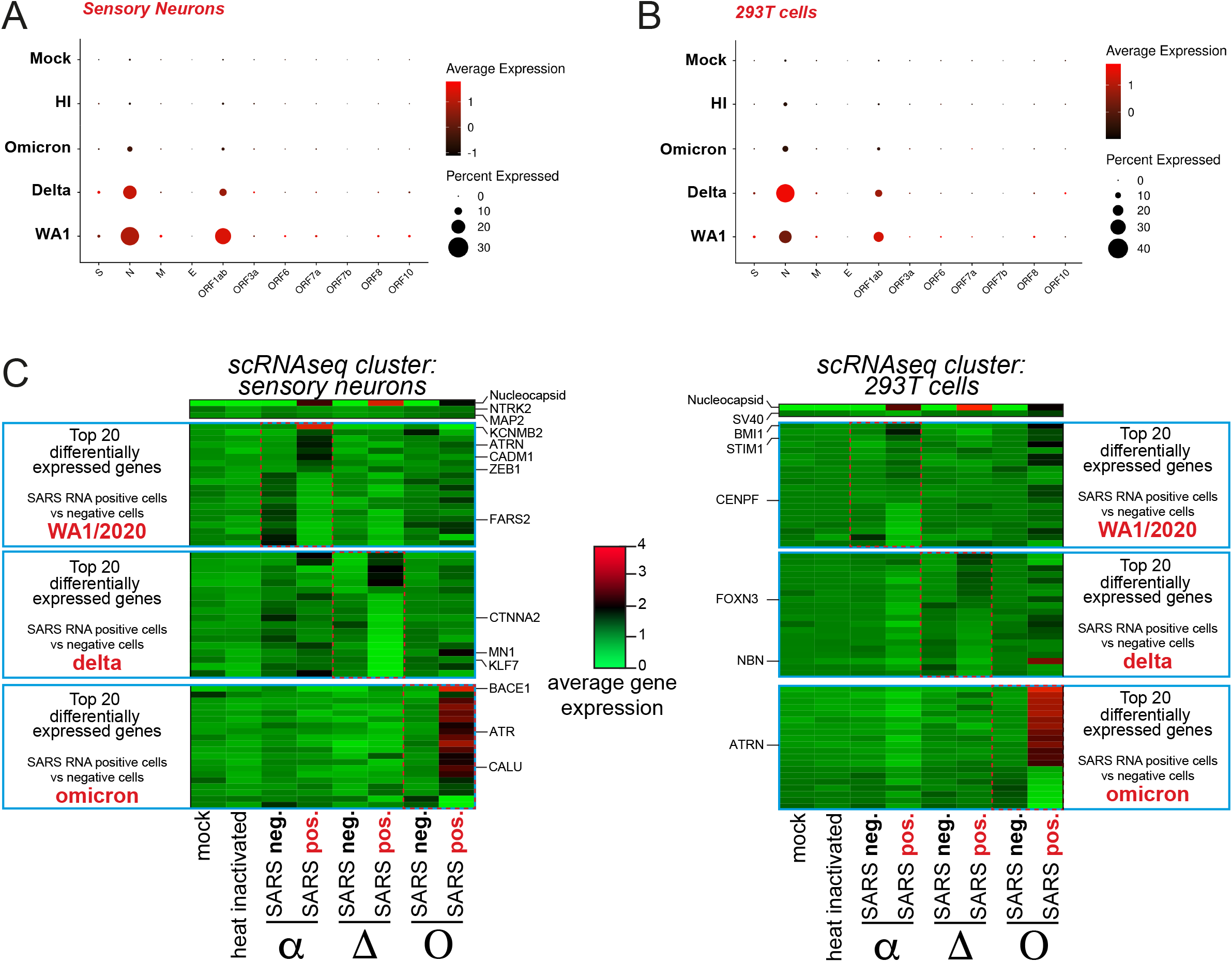
Infection of sensory neurons by SARS-CoV-2 variants delta and omicron. (A) SARS-CoV-2 genes expression in hiPSC-sensory neurons exposed to mock, heat-inactivated or active WA1/2020 SARS-CoV-2 or the delta and omicron variants. (B) SARS-CoV-2 genes expression in HEK293T cells exposed to mock, heat-inactivated or active SARS-CoV-2 strain WA1/2020 and variants delta and omicron. (C) Heatmap of average gene expression for the top 20 most differentially expressed genes between SARS-CoV-2 -positive (SARS pos.) and - negative (SARS neg.) hiPSC-sensory neurons and 293T cells upon exposure to WA1/2020 SARS-CoV-2 or the delta or omicron variants.

We performed differential gene expression analysis of SARS-CoV-2 RNA negative and positive sensory neurons and 293T cells infected with the WA1/2020 strain and delta and omicron variants. Among the top 20 differentially expressed genes we noticed a distinct gene expression pattern specific to each variant (Figure 3C). Of note, *ZEB1* was upregulated exclusively upon WA1/2020 and delta infection of sensory neurons (Figure 3C). ZEB1 is an Epithelial-Mesenchymal Transition (EMT) protein induced by SARS-CoV-2 infection which directly represses *ACE2* expression^26^. *CTNNA2* was downregulated upon WA1/2020 and delta infection of sensory neurons only (Figure 3C). CTNNA2 is a catenin protein important for actin neurofilament binding activity and crucial for neural fitness^27^. *BMI1*, coding for a polycomb group protein, was found upregulated for the three variants in 293T cells suggesting induction of senescence (Figure 3C). *STIM1*, downregulated for the delta variant but upregulated for the WA1/2020 strain and omicron variant, encodes for a transmembrane protein that has been found to promote SARS-CoV-2 infection through enhanced interferon reponse^28^ (Figure 3C). Finally, it is striking that although it infects a smaller fraction of cells than the other variants, the omicron variant induced a unique transcriptomic expression signature with some genes being strongly repressed and other activated (Figure 3C). Among those genes, *ATR* is upregulated upon omicron infection of sensory neurons (Figure 3C). *ATR* encodes for a kinase having an essential role in genome integrity. In 293T cells exposed to omicron variant, we noted the significant upregulation of *ATRN*, a gene encoding for an inflammatory molecule that is secreted to regulate chemotactic activity of chemokines (Figure 3C). Globally, the WA1/2020 strain induced a specific dysregulation of neuronal genes with genes such as *KCNMB2* and *CADM1* dysregulated in sensory neurons (Figure 3C). The delta variant downregulated neuronal transcription factor genes including *MN1* and *KLF7* in sensory neurons (Figure 3C). Finally, the omicron variant strongly upregulated genes involved in protein folding, processing and degradation such as *BACE1* and *CALU*.

In summary, these results suggest that WA1/2020, delta and omicron SARS-CoV-2 variants have the ability to infect human sensory neurons with the omicron variant being least potent for infection. Lastly, we described unique gene expression signatures specific of each variant.

## Discussion

In this study, we showed that sensory neurons of the peripheral nervous system can be infected by both the original WA1/2020 SARS-CoV-2 strain as well as the delta variant. It was thought that neurons cannot be infected by the SARS-CoV-2 due to a lack of *ACE2* expression^12^. This was demonstrated in olfactory sensory neurons and cortical neurons^12,29,30^ but recent studies showed some evidence of *ACE2* expression and presence of the virus in specific subtypes of sensory neurons. For example, SARS-CoV-2 was detected in the outer segment of photoreceptors^31^ and in the nociceptor sensory neurons of the dorsal root ganglion of COVID-19 patients^32^. Using scRNAseq, we showed that a fraction of sensory neurons was susceptible to infection (~25%). Using an iPS differentiation system, we generated a heterogenous population of sensory neurons resulting in a mixed pattern of *ACE2* expression. We showed that this population of sensory neurons can get infected however they do not release viral progeny. These *in vitro* observations with iPS-derived sensory neurons may correlate with peripheral neuropathies observed in COVID-19 patients. Sensory neurons from the dorsal root ganglion project from the lungs to the brain. The high viral load in COVID-19 patients’ lungs may stimulate infection of sensory neurons from the dorsal root ganglion.

### SARS-CoV-2 gene expression

Our results investigating SARS-CoV-2 gene expression in sensory neurons and 293T cells showed an abundance of Nucleocapsid RNA and Orf1ab RNA compared to other viral RNAs (Figure S1C and 3A-B). This observation has been made in previous studies using either cell lines or patient cells^33^. Detection of Nucleocapsid and Orf1ab RNA regions are frequently used as diagnostic tests by RT-PCR^34^. The reason why SARS-CoV-2 gene expression is biased towards these two genes is still debated. One possibility is that Orf1ab RNA, being the first part of the viral genome, is actively replicated serving for the rapid generation of RNA dependent RNA polymerase (RdRp). The abundance of Nucleocapsid RNA upon infection has also been reported and is thought to be necessary for the translation of high levels of Nucleocapsid proteins able to coat and protect newly generated SARS-CoV-2^21,33^.

### SARS-CoV-2 assembly in human sensory neurons

A striking observation is that although human sensory neurons can be infected, their infection does not produce new viruses as shown by plaque assays (Figure 2B). It is crucial to better understand the mechanism by which sensory neurons repress viral production. It has been shown that the phosphorylation landscape of host and viral proteins through CK2 and p38 MAPK kinases plays an important antiviral role^35^. Additionally, prescription of tyrosine kinases inhibitors slows down the progression of the disease^36–39^. We speculate that the repression of tyrosine kinases exclusively in infected sensory neurons may explain why SARS-CoV-2 does not get packaged and released from the cells. Furthermore, nucleocapsid protein carry an intrinsically disordered region (IDR) giving the protein the ability to phase separate with viral RNA in the cytoplasm of infected cells^40–42^. The phase separation of nucleocapsid protein with SARS-CoV-2 RNA plays an important role in the recruitment by stress granules, the inhibition of innate immunity pathways and the packaging of new viruses^42^. In host cells, nucleocapsid protein partitions with human hnRNPs such as FUS and hnRNPA2 in their phase-separated forms^41^. The formation of these robust liquid-liquid phase separations complexes composed by nucleocapsid, viral RNA and hnRNPs is key in the production of new viruses^41,42^. In our study, we show that *FUS* and two hnRNPs (*hnRNPD* and *hnRNPA2B1*) are among the top downregulated genes in infected sensory neurons versus non-infected sensory neurons (Figure 1D). We speculate that lower levels of FUS and hnRNPs in infected sensory neurons may have a negative impact on the assembly of new viruses.

## Figure legends

**Figure S1:**
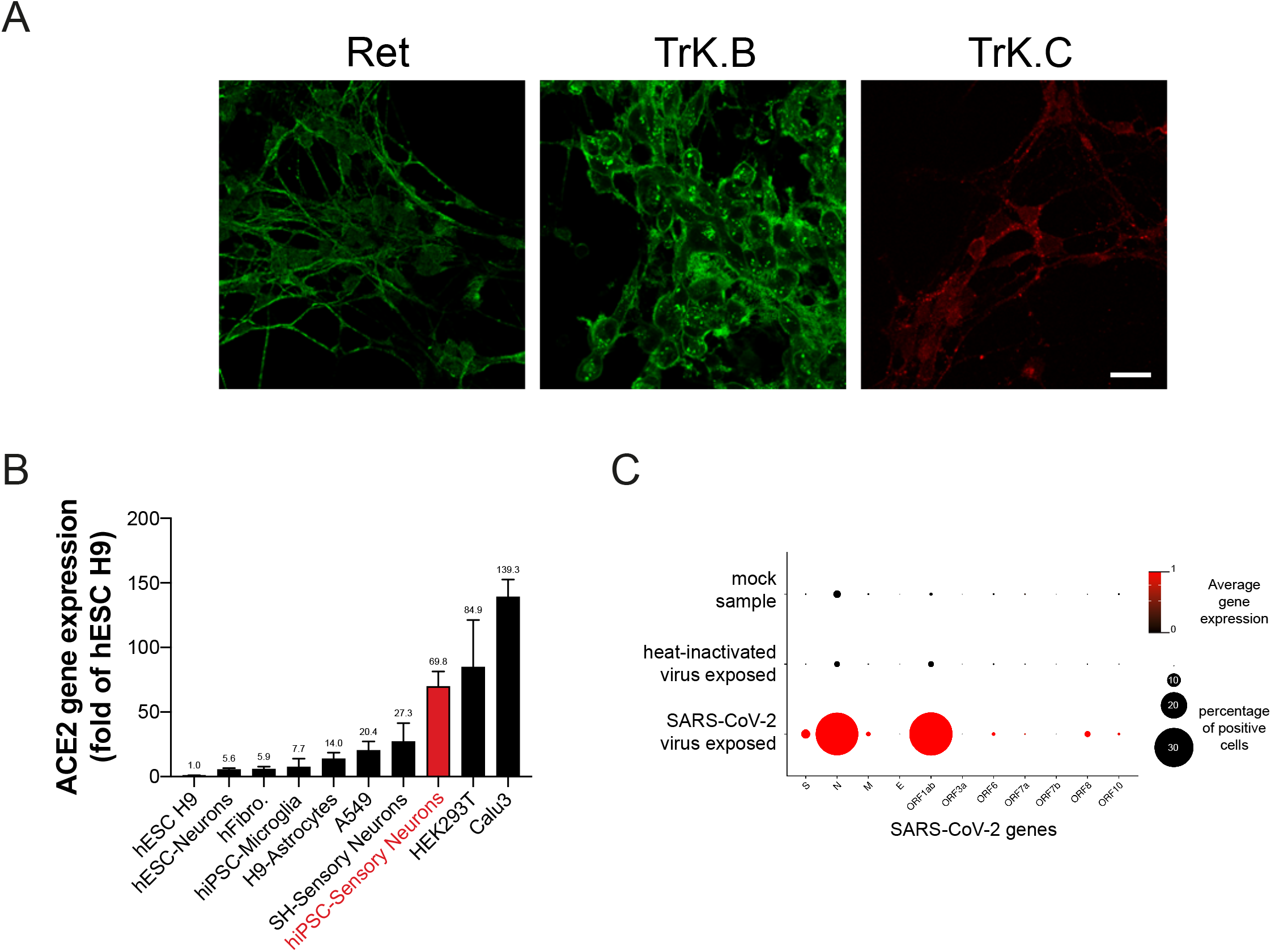
Human sensory neurons express the SARS-CoV-2 receptor gene ACE2 and can be infected by SARS-CoV-2. (A) Immunofluorescence for Ret, TrK.B and TrK.C in hiPSC-sensory neurons after 1.5 months of differentiation. Scale: 20μm (B) *ACE2* gene expression by qRT-PCR in 10 different cell types normalized to *RPLP0* housekeeping gene expression. N=3 biological replicates for each condition. Mean +/- S.E.M. (C) SARS-CoV-2 genes expression in hiPSC-sensory neurons and 293T cells co-cultures exposed to mock, heat-inactivated or active SARS-CoV-2.

**Figure S2:**
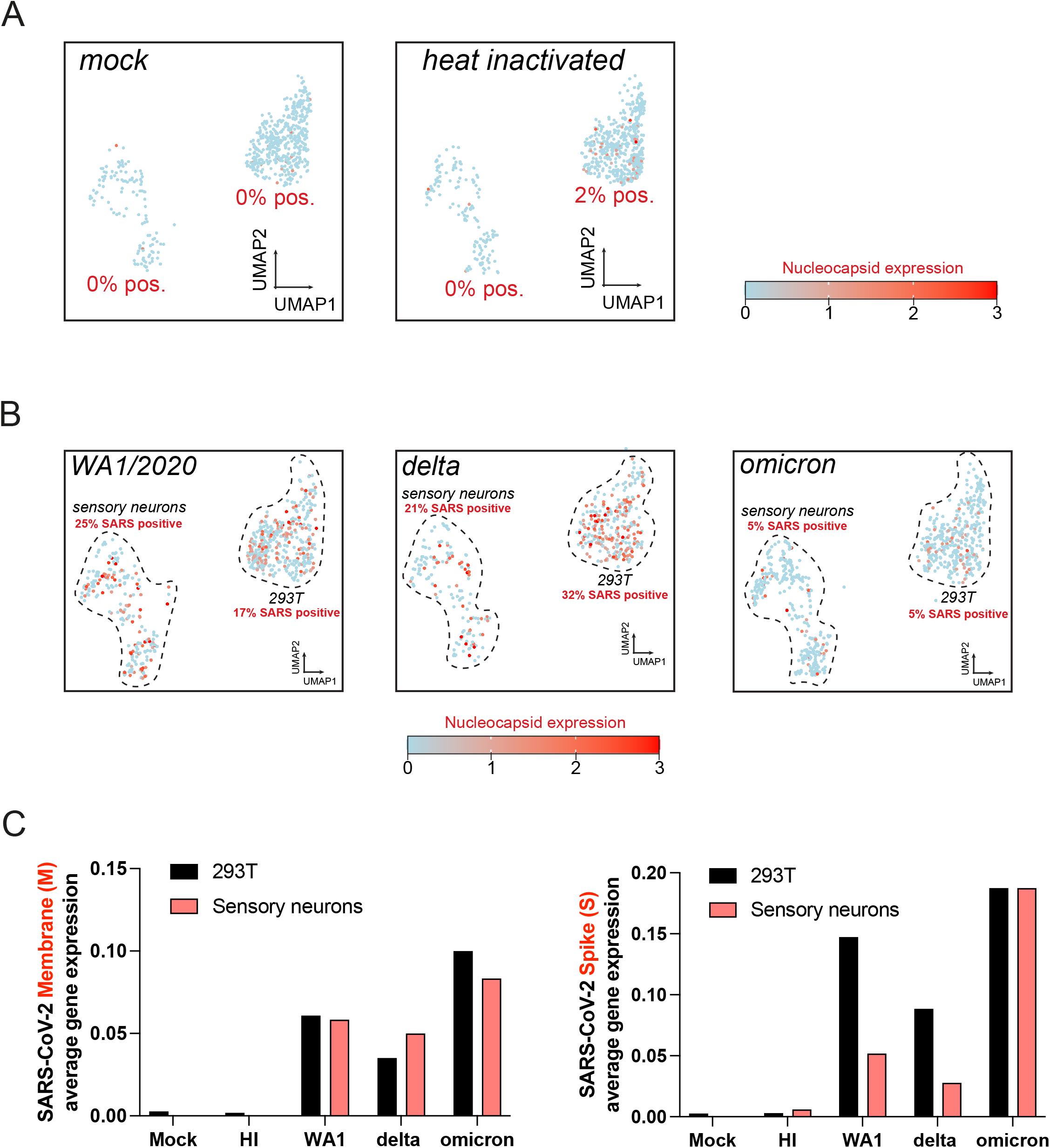
Expression of SARS-CoV-2 nucleocapsid gene by scRNAseq. (A) Feature plot of average expression level of for SARS-CoV-2 Nucleocapsid RNA in hiPSC-sensory neurons co-cultured with HEK293T cells untreated (mock) or exposed to heat-inactivated (HI) SARS-CoV-2 for 48h. (B) Feature plot of average gene expression for SARS-CoV-2 Nucleocapsid in hiPSC-sensory neurons co-cultured with HEK293T cells exposed to WA1/2020 SARS-CoV-2 or the delta or omicron variant for 48h. (C) Average gene expression for the SARS-CoV-2 Membrane (M) and Spike (S) in hiPSC-sensory neurons and HEK293T cells exposed to WA1/2020 SARS-CoV-2 or the delta or omicron variant for 48h. Mock (no virus) and heat-inactivated virus (HI) are used as controls. Data generated form scRNAseq experiments.

## Methods

### Cell culture

hiPS cell line was generated from 2-year-old male foreskin fibroblasts ordered from Coriell institute (AG07095). hiPS cell line was maintained with daily media change of mTeSR1 plus (StemCell technologies) and passaging every 4-5 days using ReLeSR (StemCell technologies). hES-qualified Matrigel (Corning) was used as a matrix. Calu3 cells and HEK293T cells were cultured using DMEM, 10% FBS, L-Glutamin, Pen/Strep and non-essential amino acids. All cells were cultured at 37°C with 5% of CO_2_.

For the co-cultures, HEK293T cells were resuspended in Neuro maturation media (BrainPhys media, N2 supplement (1X), B27 supplement (1X), BDNF (10ng/ml), GDNF (10ng/ml), NT3 (10ng/ml), Pen/Strep and non-essential amino acids) and added to terminally differentiated sensory neurons. Co-cultures were analyzed or infected after 48h at 37°C.

### hiPSC differentiation into sensory neurons

Sensory neurons from hiPSC were generated in three steps from fully pluripotent hiPSC cultures at 80% of confluency. Step 1: for the differentiation into neural crest cells (NCC), StemDiff Neural Crest Differentiation Kit (StemCell Technologies catalog #08610) was used according to manufacturer’s instructions. Step 2: NCC were expanded for at least 3 passages at a split ratio of 1:4 in NCC expansion media (DMEM/F12, N2 1X, B27 1X, non-essential amino acids, Glutamin, Pen/Strep, FGF2 (20ng/ml), EGF (20ng/ml) and CHIR99021 (2μM)). Rock inhibitor at a final concentration of 10μM was added for 24 hours after plating the cells. Step 3: sensory neuron differentiation was induced by adding neuro maturation media (BrainPhys media, N2 supplement (1X), B27 supplement (1X), BDNF (10ng/ml), GDNF (10ng/ml), NT3 (10ng/ml), Pen/Strep and non-essential amino acids) for at least 4 weeks with half media change every 3-4 days.

### Infection with SARS CoV-2

SARS-CoV-2 strain USA-WA1/2020 (WA1) was obtained from BEI Resources. SARS-CoV-2 variants B.1.617.1/Delta (Delta) and Omicron were obtained from the MassCPR variant repository. The variants were isolated at the Ragon institute BSL3 by rescue on Vero-E6 cells from primary clinical specimens. The virus stocks were prepared and tittered on Vero-E6 cells. Cells were infected with WA1, Delta and Omicron at 1 Multiplicity Of Infection (MOI) followed by incubation for 1 hour with gentle rocking every 10-15 min at 37 °C and 5% CO_2_. The medium was replaced after 1 hour with fresh medium. 48 hours post infection (48hpi), the supernatant was collected for plaque assay to determine the progeny virus production and infected cells were harvested for RNA seq assay.

### Plaque assays

Plaque assay was carried out on Vero-E6 cells in a 24-well plate format. Supernatant collected from infected cells was serially diluted in DMEM and 100 μl of the diluted sample was used to infect the monolayer of Vero-E6 cells. The plates were incubated at 37 °C under 5% CO2 for 1 hour with rocking every 10-15 min. After 1 hour, the inoculum was removed and wells overlayed with 1% methylcellulose (Thermo Fisher Scientific) in DMEM (Millipore Sigma) and plates were incubated at 37 °C up to 4-5 days, until the plaques can be observed in control wells. To visualize the plaques, the medium was removed, the cell monolayer was fixed using methanol and the cells were stained with 1% crystal violet. Plaques were counted to determine the plaque forming units per ml of media.

### Quantitative RT-PCR

RNA was extracted using RNeasy kit (Qiagen) according to manufacturer’s instruction and 1 μg of total RNA was reverse transcribed using qScript cDNA Supermix kit (Quantabio - 95048-500). 3 μl of diluted cDNA (1:20) was used with Fast SyBr green qPCR master mix (Life Technologies) for the PCR reaction. QuantStudio 6 instrument and software were used to acquire and analyze the data. Primers used for ACE2 gene expression analysis are as follow: Fwd: caagagcaaacggttgaacac; Rev: ccagagcctctcattgtagtct. Primers used for RPLP0 gene expression analysis are as follow: Fwd: gcagcatctacaaccctgaag; Rev: gcagacagacactggcaaca

### Immunofluorescence

Cells on iBidi treat 8-chamber slides were fixed for 15 min using 4% paraformaldehyde (PFA), washed one time with PBS and permeabilized 10 min using PBS/0.1% Triton X-100. Cells were blocked using PBST (PBS+ 0.1% Tween 20) with 2% Bovine Serum Albumin (BSA). Cells were incubated with primary antibodies (α-Trk.B (Rnd Systems AF1494), α-Trk.C (Rnd Systems MAB373) and α-Ret (abcam ab134100)) diluted (1:400) in PBST with 1% BSA overnight in a humidified chamber at 4°C. After 3 washes of 10 min in PBST, cells were incubated with secondary antibody (AlexaFluor – Life Technologies) diluted (1:1,000) in PBST with 1% BSA for 1 hour at room temperature. Stained cells were washed 3 times with PBST containing DAPI and mounted using iBidi mounting medium. Slides were imaged using RPI spinning disk confocal microscope.

### Single-cell RNA sequencing (scRNAseq)

scRNAseq libraries were prepared using the split-seq kit from Parse Biosciences (WT Mini V2) according to manufacturer’s instructions. A total of 20,000 cells were processed and sequenced from 2 sublibraries. Sublibraries were sequenced on a NovaSeq SP flowcell as paired-end 114×86bp for a total of 818 milions of reads (41,000 reads per cell). Fastq files were obtained with cellranger (V.4.0.0) mkfastq option. Cell-gene count matrix files were generated by Parse Biosciences pipeline using concatenated fasta sequences and GTF annotations from the human genome (ENSEMBL release 93), SARS-CoV-2 genome (MN988713.1) and SV40T sequence (NC_001669.1). To map negative strand viral reads, the strand was set to “-” in the GTF annotation file of SARS-CoV-2 genes. scRNAseq data were analyzed with the R package Seurat 4.0.3^43^.

## Acknowledgements

We thank members in the laboratories of Rudolf Jaenisch and Richard Young and other colleagues from Whitehead Institute and MIT for helpful discussions and resources. We thank Wendy Salmon from the Whitehead W.M. Keck Microscopy Facility and Whitehead Genome Core Facility. This work was supported by grants from the NIH (U19AI131135 and R01MH104610)

## Author contributions

Project design by A.F., execution of experiments by A.F., P.B., A.R. and D.T.; data analysis by A.F., A.R. and D.T.; manuscript preparation by A.F., P.B., A.R., D.T. and R.J.

